# Strength of enemy release from parasitoids is context-dependent in the invasive African Fig Fly, *Zaprionus indianus*

**DOI:** 10.1101/2024.07.09.602257

**Authors:** Camille R. Walsh-Antzak, Priscilla A. Erickson

## Abstract

Understanding the mechanisms underlying the success of biological invasions is essential to employ effective prediction and management strategies. Escape from natural enemies in invaded regions (enemy release hypothesis, ERH) and increased competitive ability are hallmarks of invasive species; however, these two processes are rarely studied within the same context. Here, we examined the effect of enemy release on the competition outcomes of a successful invasive insect pest in North America, the African fig fly (*Zaprionus indianus)*. Parasitoid wasps such as *Leptopilina heterotoma* that parasitize drosophilid larvae may seek out established species with known host suitability over a novel species, so we hypothesized *Z. indianus* may have low susceptibility to parasitoids, giving them a competitive advantage over co-occurring drosophilids. We tested this hypothesis by comparing the adult emergence rates from *Z. indianus* larvae reared alone or in competition with *Drosophila hydei* or *D. simulans* larvae in the presence and absence of parasitoid wasps under low and high larval densities. At low larval densities, *Z. indianus* emerged at equal rates to *D. hydei* but outcompeted *D. simulans*, and these outcomes were not affected by parasitoids. However, at high densities, the addition of parasitoids shifted competition outcomes in favor of *Z. indianus*, suggesting enemy release provides a competitive advantage under some circumstances. These results indicate that the strength of enemy release in *Z. indianus* is widely dependent on contextual factors such as density and competitor species. This study emphasizes how a community approach to testing the ERH is vital as the overall interpretation of the presence and strength of enemy release differed between intraspecific and interspecific experiments. Further investigation of how these results apply to field environments could offer insight into how *Z. indianus* alters ecosystems and how productive biological control may limit the spread of *Z. indianus*.

**Short Abstract:** Invasive species may succeed in new environments in part because they are less susceptible to diseases and parasites that have co-evolved with local hosts, giving invaders a competitive advantage. We tested this hypothesis by competing an invasive fruit fly against established species in the presence of parasitoid wasps that lay their eggs in fruit fly larvae. We found that the invasive species generally outcompeted other species in the presence of parasitoids, but the extent of its advantage depended on the species it was competing against and the number of larvae present.

## Introduction

The spread of invasive species poses significant ecological, environmental, and economic threats. Invasive species have largely contributed to biodiversity loss and the extinction of native species (Bradshaw et al. 2016) and are destructive in novel environments, leading to agricultural damage and economic costs (Diagne et al. 2021). Furthermore, invasive species raise human health concerns as they can alter disease-vector dynamics and facilitate the spread of pathogens (Roy et al. 2022). Despite the well-known risks of biological invasions, managing invasive species remains a challenge. A comprehensive understanding of the mechanisms underlying invasive species’ success is vital for effective conservation, management, and mitigation practices.

One of the most widely proposed hypotheses explaining the success of invasive species is the enemy release hypothesis (Keane & Crawley 2002, Enders et al. 2018). This hypothesis proposes that invasive species in novel environments are no longer impacted by their natural enemies (e.g., predators, parasites, or pathogens from the invader’s native range). As a result the invader population grows as local species, but not the invader, are restrained by natural enemies. Due to the loss of natural enemies, invasive species may also succeed in invaded communities by relocating resources from defence mechanisms to growth and development (evolution of increased competitive ability hypothesis, Blossey & Notzold 1995). Competition with native species also plays a crucial role in determining the success or failure of invasive species (Clarke & McGeoch 2023). The inherent competitive ability of invasive species is often superior to native species, facilitating their dominance in invaded communities (Gioria & Osborne 2014, Fridley & Sax 2014). Although components of enemy release and competition overlap, there are few studies examining both as factors for invasion success. Studying these two processes within the same context may provide a more cohesive understanding of the spread and success of invasive species.

The African fig fly, *Zaprionus indianus*, is a member of the family Drosophilidae and is native to tropical regions of central Africa. *Z. indianus* is a polyphagous consumer of decaying fruit and has over 80 documented host species, though it can also be a pest on ripening figs (Pfeiffer et al. 2019) and potentially raspberries (Stazzonelli et al. 2023). In the 1990s, *Z. indianus* was first detected in South America in Brazil, where it quickly spread across the continent and damaged fig crops (Vilela 1999). Following a northward invasion pattern, *Z. indianus* was detected in Florida in 2005 (Van der Linde et al. 2006) and has since been detected across eastern North America (Pfeiffer et al. 2019, Rakes et al. 2023). Sampling data from apple and peach orchards reveal repeated and rapid population growth relative to other drosophilids, suggesting *Z. indianus* outcompetes co-occurring drosophilid species (Rakes et al. 2023).

However, the factors that facilitate the success of *Z. indianus* remain unknown. Evaluating the role of enemy release in competition outcomes could shed light on the mechanisms underlying the repeated success of *Z. indianu*s in novel environments. *Zaprionus indianus* is an optimal study system as its relatively recent arrival to North America and lack of permanent establishment in temperate regions (Pfeiffer et al. 2019) suggests it has not co-evolved with North American enemies, providing an opportunity to assess whether reduced enemy pressure in a novel environment mediates their population success.

Parasitoid wasps are natural enemies of drosophilids and exert strong selective pressures on fly populations, with up to 90% of drosophilids parasitized by parasitoids in certain environments (Fleury et al. 2004). Additionally, host-parasitoid interactions exhibit coevolutionary arms races, and each have highly specialized attack and defense mechanisms (Godfray & Shimada 1999). Invasive species lack this coevolutionary relationship with parasitoids present in the introduced range which raises questions about how they interact with these novel parasitoids (Kraaijeveld et al. 1998). *Leptopilina heterotoma* is a generalist parasitoid wasp found in North America that parasitizes larvae of numerous drosophilid species due to its low selection threshold (Quicray et al. 2023, de Bruijn et al. 2022). The life cycle of *L. heterotoma* begins with an adult parasitoid ovipositing into fly larvae. If the parasitization is successful (e.g., the parasitoid overcomes the flies’ immune defense), then the parasitoid develops inside the host and slowly consumes it, ultimately resulting in host death during fly pupation. An adult parasitoid typically emerges from the fly pupae ∼21-23 days after oviposition (Quicray et al. 2023). Although *L. heterotoma* is a generalist, not all hosts are suitable for parasitoid development (Quicray et al. 2023). While *L. heterotoma* detects *Z. indianus* as potential hosts (e.g., ovipositing in larvae), the majority of *Z. indianus* successfully emerge as adults, suggesting that the immune system of *Z. indianus* is resistant to *L. heterotoma*, perhaps due to the presence of giant hemocytes (Kacsoh et al. 2014). Additionally, no *L. heterotoma* adults emerge from *Z. indianus*, indicating *Z. indianus* is an unsuitable host. Therefore, invasive populations of *Z. indianus* might benefit relative to other drosophilids by escaping parasitism from *L. heterotoma*.

Resource competition is another major driving force behind the success and spread of invasive species as they often have greater capacities to exploit resources more efficiently than local species (Mooney & Cleland 2001, Gioria & Osborne 2014). Ephemeral food substrates, such as rotting fruits and their associated microbiota, result in both intra- and interspecific competition for drosophilids (Rohlfs & Hoffmeister 2004). *Z. indianus* likely experiences high rates of resource competition with other drosophilid species because it occupies similar niches (e.g., opportunistic oviposition in rotting host fruits) (Bragard et al. 2022). Competition outcomes in drosophilids are widely context-dependent and may fluctuate based on abiotic and biotic factors such as air temperature or fruit substrate (Comeault & Matute 2021, de Paiva Mendonça 2023). Host-parasitoid interactions can also influence competition outcomes. For example, *D. melanogaster* outperforms its closest relative, *D. simulans*, in the absence of parasitoids, but these competition outcomes are inverted when parasitoids are present (Boulêtreau et al. 1991). However, it is unknown how parasitoids may alter competition outcomes between recent invaders and local populations. If *L. heterotoma* preferentially parasitizes established species in the presence of *Z. indianus*, then *Z. indianus* may capitalize on reduced competition stressors.

In this study, we evaluated how parasitoids affect the competition outcomes of *Z. indianus* under controlled laboratory conditions. As our main goal was to evaluate how enemy release influences the dominance of *Z. indianus* in invaded environments, we used a community-comparison approach as described by Brian & Catford (2023). We measured adult emergence rates in the absence and presence of *L. heterotoma* to compare competition outcomes both between and within species. We conducted assays at low density and high density treatments to account for varying fly densities observed in the field across different locations and time periods, which may provide insight about host-parasitoid interactions in spatially and temporally heterogeneous environments (Rakes et al. 2023, Gleason et al. 2019). In the interspecific competition assays, we offered parasitoids the opportunity to parasitize the larvae of *Z. indianus* and another drosophilid species. We used *D. simulans* and *D. hydei* as co-occurring, established species as they are both cosmopolitan in distribution (Werner et al. 2018). Although not native to North America (Zhao & Begun 2017, Lachaise et al. 1988), *D. simulans* and *D. hydei* have both been widespread in the eastern US since at least 1921 (Sturtevant 1921), allowing substantially more time for coevolution with North American parasitoids relative to *Z. indianus*. Genomic data suggest that despite a population bottleneck, introduced North American *Z. indianus* populations retain substantial genetic variation relative to congeners (Comeault et al. 2020, 2021). Therefore, the strength of enemy release may vary due to genetic variation. We compared the parasitized emergence rates of a low-latitude population from Florida with a high-latitude population from Connecticut, as flies from Florida and temperate regions are genetically differentiated (Erickson et al. 2024). Lastly, we evaluated emergence rates with differing parasitoid exposure times to test whether *L. heterotoma* exhibits host-switching (Murdoch 1969), in which a parasitoid initially oviposits in a favorable host species, but then switches to a different host after the original host becomes saturated.

Given the lack of coevolutionary history and previously described host incompatibility of *L. heterotoma* to *Z. indianus*, we hypothesized that *Z. indianus* would be less susceptible to parasitization than established drosophilids, leading to higher emergence rates for *Z. indianus*. Given the rapid population growth of *Z. indianus* observed in orchards (Rakes et al. 2023), we predicted that *Z. indianus* performs equal to or better than established species in the absence of parasitoids, and this advantage is further intensified by the presence of *L. heterotoma*, alleviating competition stressors for *Z. indianus*.

## Materials & Methods

### Insect rearing

Adult *Z. indianus* were obtained from isofemale lines derived from females collected in North America in 2022 and 2023 (Rakes et al. 2023). *D. simulans* were collected from orchards in Florida in 2022, and *D. hydei* were collected from orchards in Virginia in 2023. All flies were maintained on a cornmeal-molasses artificial diet at room temperature. All parents of the experimental offspring were raised in controlled lab conditions for at least two generations. *L. heterotoma* (Lh14) stocks were obtained from Todd Schlenke at the University of Arizona. The Lh14 strain was originally collected in 2002 from Winters, California (Kacsoh & Schlenke 2012). Parasitoids were maintained on laboratory strains of *D. melanogaster* (w1118 and Canton-S). To elongate parasitoid lifespan, 500 μL of 50% honey-water solution was provided on vial plugs.

### Intraspecific competition assays

To test how parasitoid presence affects intraspecific competition outcomes of *Z. indianus*, we prepared vials with only *Z. indianus* larvae in the absence and presence of parasitoid wasps. We allowed adult flies to oviposit on 3% agar plates containing 10% grape juice concentrate for 24 hours. A paste of baker’s yeast and water was added to each plate to encourage oviposition. We collected two-day-old larvae from the plates and inoculated vials filled with 10 mL of cornmeal-molasses substrate. We compared emergence rates at low densities (50 total larvae) and high densities (150 total larvae). For parasitized treatments, we immediately added six adult *L. heterotoma* (three female, three male) to the inoculated vials for 24 hours. Parasitoids were not offered hosts prior to experiments. In the control treatments, we followed the same procedure but did not add parasitoids. We counted emerging adult flies every 1-2 days until no further adults emerged. Intraspecific assays had two replicates per treatment. We conducted two independent intraspecific assays (one alongside each interspecific assay, see below) and combined the data for a total of four vials per density and parasitoid treatment.

### Interspecific competition assay

To test how parasitoid presence affects interspecific competition outcomes, we compared the larva-to-adult survival rates of *Z. indianus* and co-occurring drosophilid species (*D. hydei or D. simulans)* reared together in the absence and presence of parasitoid wasps. Low and high total densities were the same as the intraspecific assays (50 and 150 larvae); however, interspecific assays contained equal numbers of each species (25 larvae of each species for the low density experiment and 75 of each species for high density). We collected larvae as described above from the two species separately and combined them into a single vial with or without parasitoids. We added six parasitoids (three female, three male) for 24 hours. We counted adults of both species every 1-2 days as they emerged. Each interspecific treatment had five replicates, totaling 20 vials per species comparison. All experiments were incubated at room temperature.

### Comparing parasitoid susceptibility across Z. indianus populations

To test for potential genetic differences in parasitoid susceptibility, we compared emergence rates of *Z. indianus* from five Florida (FL) isofemale lines, and four Connecticut (CT) isofemale lines collected in 2022 (Rakes et al. 2023). These isofemale lines were reared in the lab for approximately 16 months (∼20 generations) prior to the experiments described here. We added a total of 25 *Z. indianus* larvae to each vial. We prepared three replicates of parasitized vials for each line, resulting in a total of 27 vials. Two parasitoids (one female, one male) were added to vials for 24 hours.

### Assessing potential parasitoid host-switching

As *D. simulans* is a known suitable host of *L. heterotoma* (Quicray et al. 2023), we predicted that *L. heterotoma* might initially oviposit in *D. simulans* before parasitizing *Z. indianus*. To assess parasitoid foraging behavior, we compared the emergence rates of *Z. indianus* and *D. simulans* following 1 hr, 8 hr, and 24 hr of parasitoid exposure. All vials contained a low density of 20 larvae (10 of each species) to increase the likelihood of host saturation. Two parasitoids (one female, one male) were added to parasitized treatments. Non-parasitized treatments experienced 0 hr of parasitoid exposure. Each parasitoid duration treatment had three replicates (9 total vials), and the non-parasitized controls had 12 replicates.

### Data Analysis

All analyses were performed using the R statistical software (v. 4.4.0; R Core Team 2024). We used the proportion of adult flies relative to starting larvae to calculate emergence rates of each treatment. We used a generalized linear model (GLM) with a binomial distribution to compare *Z. indianus* emergence rates for the intraspecific assays. Emerged adult flies were modeled as a function of *density* (as a factor), *parasitoid presence*, and a *density*parasitoid* interaction. We analyzed the interspecific data in separate models for each density and competing species comparison. We used GLMs with a quasibinomial distribution to compare species emergence rates between treatments. The quasibinomial distribution corrected for overdispersion in the interspecific data (variance/mean ratios ranged from 5-10). Emerged adult flies were modeled as a function of *species, parasitoid presence*, and a *species*parasitoid* interaction. We used a generalized linear mixed model (GLMM) implemented in *lme4* (Bates et al. 2015) with a binomial distribution to compare *Z. indianus* emergence rates from Florida (FL) and Connecticut (CT) populations. The model included *location* as a main effect and *isofemale line* as a random effect. We used a GLM with a binomial distribution to assess differences in *Z. indianus* and *D. simulans* emergence rates at various parasitoid exposure times as a function of the *species, exposure time*, and a *species*exposure time* interaction.

We calculated estimated marginal means (least squared means) for each model and performed pairwise linear contrasts between groups using the *emmeans* package (Lenth et al. 2020) in R. We focused on comparing groups that differed by only a single variable (e.g. same species but different parasitoid treatment, or same treatment but different species, Appendix Table 1). We generated Tukey-adjusted linear contrast P values; statistical significance was determined by P_adjusted_ < 0.05. All plots were generated with *ggplot* (Wickham 2016) and data were managed with *data*.*table* (Dowle & Srinivasan 2012).

## Results

### Intraspecific competition assay

High density (GLM; Z = -8.326, P < 0.001) and presence of *L. heterotoma* (Z = -8.600, P < 0.001) significantly decreased *Z. indianus* emergence rates (Fig. 1A, Appendix Table 1). There was a significant interaction between parasitoid presence and density, with parasitoids causing a greater reduction in emerging adults at low density relative to high density (Z = 4.729, P < 0.001).

**Figure 1:**
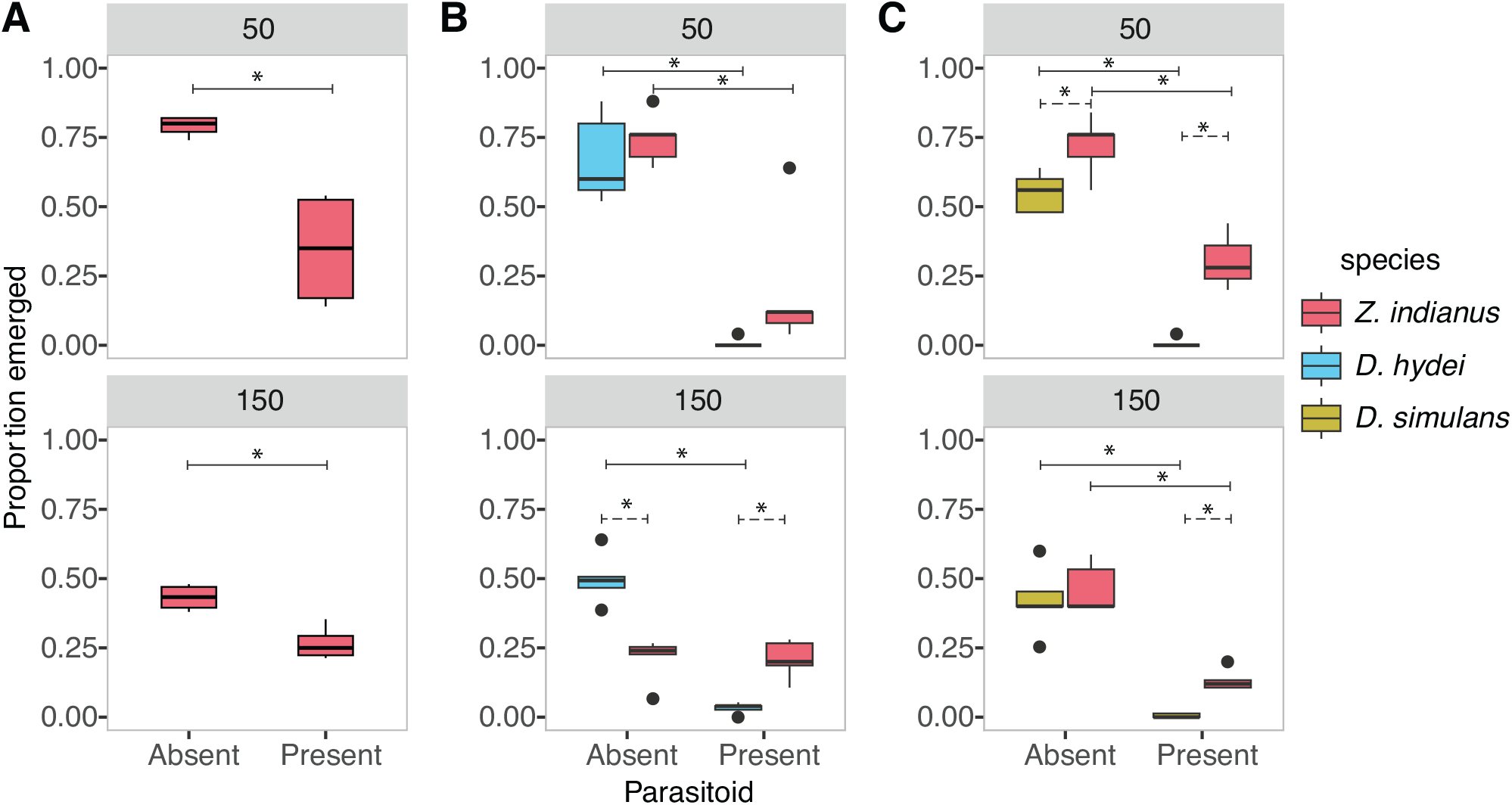
Enemy release from *L. heterotoma* depends on competing species and larval density in *Z. indianus*. A) Intraspecific competition assay results comparing emergence rates of unparasitized and parasitized *Z. indianus* larvae. Gray labels indicate density as the total number of larvae present. B) Interspecific competition assay results comparing *Z. indianus* and *D. hydei* and C) *Z. indianus* and *D. simulans* adult emergence rates. The low density (top) is 50 total larvae (25 of each species in B-C) and the high density (bottom) is 150 total larvae (75 of each species in B-C). Asterisks indicate statistical significance by linear contrast of a generalized linear model (P_adjusted_ < 0.05). Solid lines with asterisks indicate significant effects of parasitoids within species. Dotted lines with asterisks indicate significant differences between species raised in direct competition. N=5 replicate vials per treatment.

### Interspecific competition assay

At low density, parasitoid presence significantly decreased the emergence rates of co-reared *Z. indianus* and *D. hydei* (linear contrast of GLM; Z = 4.234, P < 0.001 & Z = 2.837, P = 0.024, respectively; Fig. 1B top panel, Appendix Table 1). However, there were no significant differences between *Z. indianus* and *D. hydei* emergence rates in either the absence or presence of parasitoid wasps (Z = -0.654, P = 0.914 & Z = -1.748, P = 0.299, respectively). At high density, the presence of parasitoids significantly decreased the emergence rates of *D. hydei* (Z = 7.402, P < 0.001; Fig. 1B bottom panel), but did not affect *Z. indianus* emergence rates (Z = 0.061, P = 0.999). *Z. indianus* had significantly lower emergence rates than *D. hydei* in the absence of parasitoids (Z = 5.445, P < 0.001), but had significantly higher emergence rates in the presence of parasitoids (Z = -4.383, P < 0.001).

At low density, parasitoid presence significantly decreased the emergence rates of co-reared *Z. indianus* and *D. simulans* (Z = 6.360, P < 0.001 & Z = 4.924, P < 0.001, respectively; Fig. 1C top panel, Appendix Table 1). However, *Z. indianus* emergence rates were significantly higher than those of *D. simulans* in both the absence and presence of parasitoids (Z = -2.737, P = 0.032 & Z = -3.898, P < 0.001). At high density, parasitoid presence also significantly decreased the emergence rates of both *Z. indianus* and *D. simulans* (Z = 6.288, P < 0.001 & Z = 4.586, P < 0.001, respectively; Fig. 1C bottom panel). There were no significant differences between *Z. indianus* and *D. simulans* emergence rates in the absence of parasitoids (Z = -0.787, P = 0.861). However, *Z. indianus* had significantly higher emergence rates than *D. simulans* in the presence of parasitoids (Z = -3.098, P = 0.011).

### Comparing parasitoid susceptibility across Z. indianus populations

There were no significant differences in adult emergence rates between parasitized Connecticut and Florida populations (Fig. 2; Appendix Table 1; GLMM; P = 0.561).

**Figure 2:**
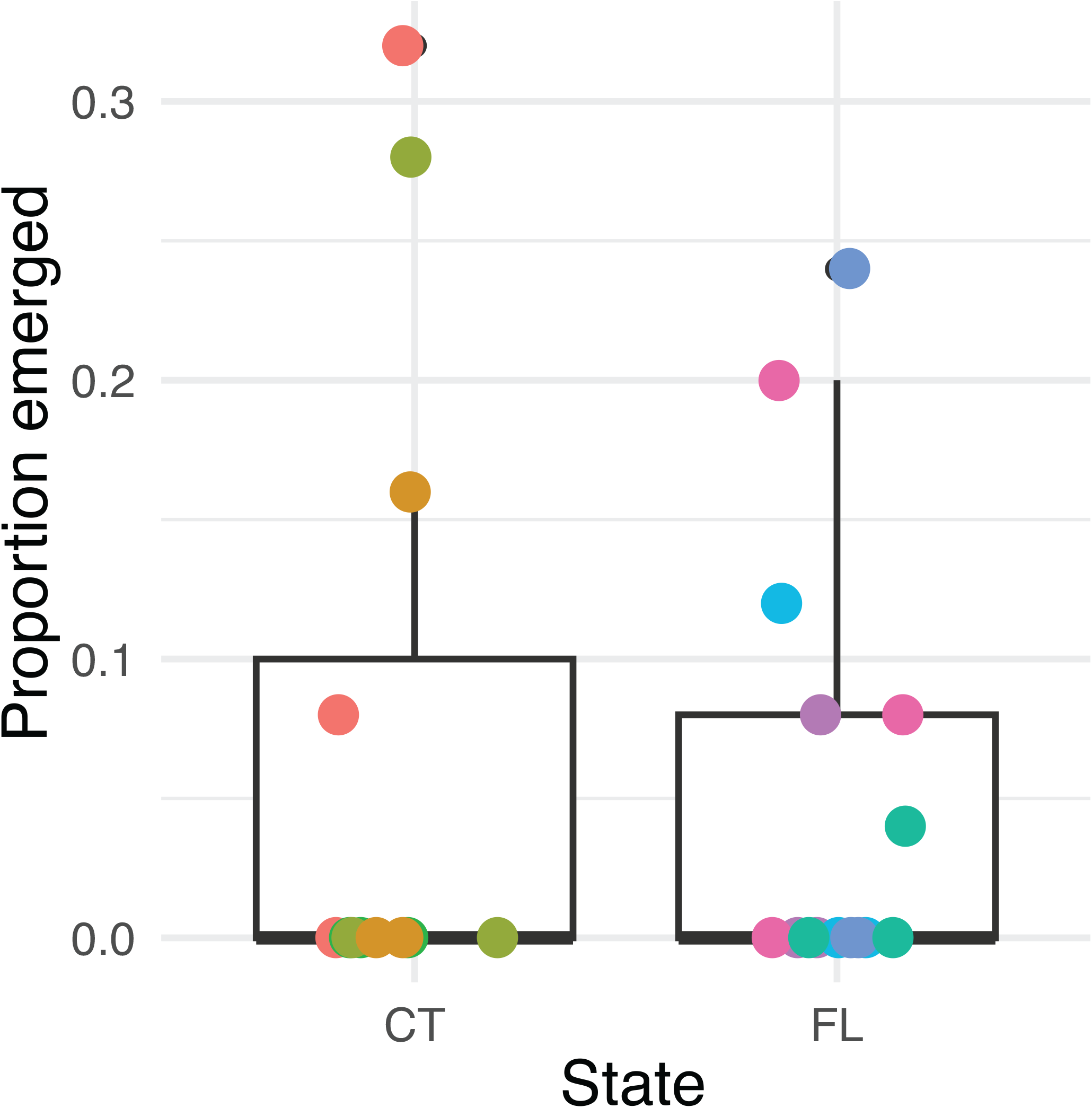
No geographic variation in *Z. indianus* susceptibility to *L. heterotoma*. Adult emergence rates from parasitized Connecticut (CT) and Florida (FL) *Z. indianus* isofemale lines. Boxplots indicate combined emergence rates of all lines and replicates from each state. Points (with horizontal jitter added for clarity) indicate individual replicate vials; colors indicate unique isofemale lines from CT (N =4 lines) or FL (N=5 lines) with three replicates of 25 larvae per line.

### Host switching assay

Exposure to parasitoids for one hour significantly decreased the emergence rates of *D. simulans* (Z = -3.819, P < 0.001), but not *Z. indianus* (Z = -1.560, P = 0.119), when compared to non-parasitized controls of each species (Fig. 3; Appendix Table 1). However, exposure to parasitoids for 8 hours and 24 hours significantly decreased the emergence rates of both *D. simulans* (P8h < 0.001, P24h < 0.001) and *Z. indianus* (P8h < 0.001, P24h = 0.004).

**Figure 3:**
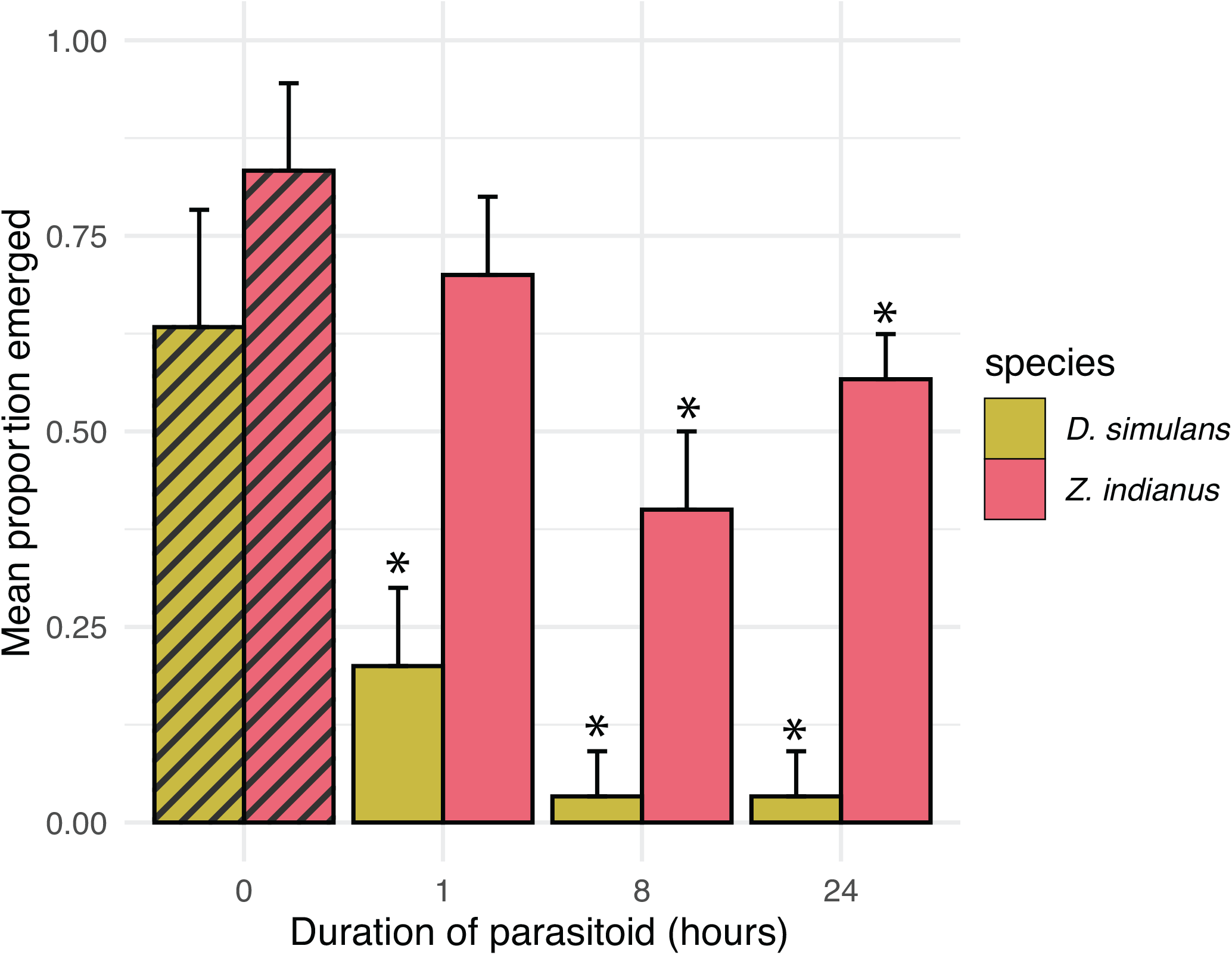
*L. heterotoma* parasitizes *D. simulans* more rapidly than it parasitizes *Z. indianus*. Interspecific assay comparing the emergence rates of *Z. indianus* (red bars) and *D. simulans* (yellow bars) at different parasitoid exposure times. Hashed lines represent control vials with no parasitoids added (N=12 total). Parasitized treatments had three replicates for each exposure time. A density of 20 larvae (10 of each species) was used for all vials. Error bars represent standard deviation. Asterisks indicate significant differences between control and parasitized treatments by linear contrast (P_adjusted_ < 0.05).

## Discussion

We explored the effect of enemy release on influencing competition outcomes by comparing the adult emergence rates of a recent invasive insect pest, *Z. indianu*s, and co-occurring cosmopolitan drosophilid species in the absence and presence of *L. heterotoma* parasitoid wasps. Here, we demonstrate that consideration of competition interactions and environmental context is essential when assessing the relative strength of enemy release. Our results suggest that enemy release in the presence of competing species may contribute to the rapid population growth of *Z. indianus* that is commonly observed in the field, and drosophilid density affects the relative strength of enemy release in benefiting *Z. indianus* populations. We found that under most circumstances, *Z. indianus* is an equal or better competitor than *D. hydei* and *D. simulans*, but competition outcomes are widely dependent on factors such as parasitoid presence, density, and competitor species. Our findings support the enemy release hypothesis because the emergence rates of *Z. indianus* were equal to or higher than *D. hydei* and *D. simulans* in the presence of parasitoids (Fig. 1B-C), indicating that parasitoids were often more detrimental to the established species (a known host) than to *Z. indianus* (a novel host), especially at high densities. Further, our results demonstrate the importance of taking a community approach when studying enemy release because *Z. indianus* was susceptible to parasitization by *L. heterotoma*, but the relative impact of this susceptibility depended on other contextual factors.

We observed that the relative strength of enemy release differed between densities, suggesting the benefit of enemy release to *Z. indianus* is density-dependent. At low densities, the addition of parasitoids did not alter competition outcomes for *Z. indianus* with *D. hydei* or *D. simulans*. Therefore, at low densities, the benefit of enemy release is small for *Z. indianus* as parasitoids have approximately equal effects on both competing species and do not alter competition dynamics. However, at high densities, we observed that parasitoid presence shifted the competition outcomes in favor of *Z. indianus* in both interspecific assays. Since competition outcomes switched to benefit *Z. indianus* at high densities, but not low densities, we conclude that enemy release was stronger at high densities compared to low densities.

A possible explanation for differing competition outcomes based on density is that *L. heterotoma* switched from a known host (D. *simulans* or *D. hydei*) to a novel host (*Z. indianus)*. As host switching is likely initiated by optimal foraging behavior (Křivan 1996), an increase in the abundance of suitable hosts (e.g., high density) might decrease the number of parasitized suboptimal hosts. We tested the possibility of host switching by examining the effects of parasitoid exposure time on the emergence rates of *Z. indianus* and *D. simulans* (Fig. 3). Our data suggest parasitoids initially oviposited in *D. simulans*, but switched to *Z. indianus* once suitable hosts were saturated. The difference in parasitization rates might result from variations in behavioral avoidance strategies between *Z. indianus* and *D. simulans* larvae, or *L. heterotoma* may prefer to oviposit in *D. simulans*. Although we did not directly assess parasitoid host preference, parasitoids are highly sensitive to host-selection and can accept or reject hosts based on their suitability and quality (Wertheim 2022). *D. simulans* primarily uses lamellocytes for its anti-parasitoid immune response, rather than multinucleated giant hemocytes like *Z. indianus* (Salazar-Jaramillo et al. 2014, Cinege et al. 2020). Virulence factors of *L. heterotoma* have evolved mechanisms to combat the lamellocyte immune response (McGonigle et al. 2017), suggesting that species using lamellocytes may be more favorable hosts than species using multinucleated giant hemocytes. Therefore, it is possible that the density-dependent results observed in the interspecific assays reflect host switching behavior of *L. heterotoma*. Although host-switching is commonly observed in generalist parasitoids (Murdoch 1969, Jones et al. 2015), specialist parasitoid species may be more reluctant to oviposit in suboptimal hosts due to increased host selectivity. Therefore, evaluation of how different parasitoid species interact with *Z. indianus* alongside other drosophilids would be beneficial for a broader understanding of enemy release since parasitoid life-history strategies may produce different competition outcomes.

Additionally, the strength of enemy release differed based on the competing species. At high densities, the strength of enemy release was greatest when *Z. indianus* was competing with *D. hydei* as the presence of parasitoids shifted competition outcomes from *Z. indianus* having significantly lower emergence rates to significantly higher emergence rates than *D. hydei* (Fig. 1B). Although parasitoid presence also shifted competition outcomes in favor of *Z. indianus* when competing with *D. simulans*, the magnitude of the shift was smaller as *Z. indianus* already performed equal to *D. simulans* in the absence of parasitoids. Although the presence of parasitoids decreased *D. simulans* and *D. hydei* survivorship to zero, parasitized larvae likely still competed with *Z. indianus* larvae as host death does not occur until after the fly has pupated (Quicray et al. 2023). We did not directly test for the presence of competition, and the density-dependent effects on survival we observed may be due to other factors such as accumulation of waste. However, a similar study directly tested for interspecific competition between *Z. indianus* and *D. suzukii*, suggesting competition is present between other species (de Paiva Mendonça 2023). Further evaluation of how competing species’ traits may alter the strength of enemy release would be beneficial for identifying the specific mechanisms that influence enemy release strength in community environments. These nuances emphasize how using a community approach for testing the enemy release hypothesis is crucial as interpretation of the strength of enemy release may differ based on factors such as density and competing species. Our results extend the work of Brian & Catford (2023) as we found that enemy diversity and the relative enemy impact can independently contribute to the strength of enemy release.

Interestingly, we observed that parasitoids significantly decreased the emergence rates of *Z. indianus* in intraspecific assays (Fig. 1), and *Z. indianus* emergence rates were never higher than 50% in the presence of parasitoids. Our *Z. indianu*s results differ from a study by Kacsoh et al. (2014), which reported the majority (> 50%) of *Z. indianus* escaped parasitism by *L. heterotoma*, but they are similar to a study by Cinege et al. (2023) which reported fewer than 30% of *Z. indianus* emerged in the presence of *L. heterotoma*. It is unlikely that genetic variation between *Z. indianus* lines is responsible for these differences, as we found no reproducible variation in emergence rates across isofemale lines (Fig. 2). Differences in laboratory conditions or insect rearing may have produced different parasitization rates across studies. Within our experiments, there were more vials with 0% emergence in the geographic variation experiment (Fig. 2) compared to the low density treatment of the intraspecific experiments (Fig. 1A). It is possible that a lower density of flies in the geographic variation experiment (N=25) led to a faster saturation rate of larvae (Mitsui & Kimura, 2000). Alternatively, an uncontrolled experimental factor, such as parasitoid health or age, may have affected parasitization efficiency between experiments.

All experiments in this study were conducted under controlled laboratory conditions, which limits applicability to field environments. We used larva-to-adult emergence rates as an indicator for population success and did not directly measure parasitization, but our use of control treatments allows us to infer that parasitization occurred. Additionally, we did not test other fitness components such as growth, fecundity, and reproductive output, which may also be used to assess enemy impact. Assessment of multigenerational effects using multiple fitness factors may more comprehensively determine the impact of parasitoid presence on invader population performance. In *D. melanogaster*, adult females that successfully defend themselves from parasitism have several reduced fitness traits such as smaller body size and lower fecundity (Fellowes et al. 1999). Additionally, investment in parasitoid defense mechanisms (e.g., increased hemocyte counts) is associated with tradeoffs in competitive ability in *D. melanogaster* (Kraaijeveld et al. 2001). These tradeoffs might particularly affect *Z. indianus*, because of their higher resistance to parasitoids due to the role of multinucleated giant hemocytes in their immune response (Kacsoh et al. 2014, Cinege et al. 2023). Therefore, increased defense mechanisms may lead to diminished fitness outcomes for *Z. indianus*, which could contribute to drops in wild population sizes seen later in the season (Rakes et al. 2023). In addition to physiological defense mechanisms, behavioral responses are also vital for reducing parasitism in hosts (Hart 1990). Assessing whether the presence of parasitoids or competing species can influence *Z. indianus’* oviposition or larval foraging behavior may provide insights into potential behavioral resistance strategies of *Z. indianus* in field environments. Further evaluation of how parasitism affects *Z. indianus* population dynamics over time would be beneficial to understand the strength and frequency of enemy release in the field. Additionally, using a more natural experimental design (e.g., natural fruit substrates, multiple co-occurring drosophilid species) to mirror environmental heterogeneity could provide more information on how these results apply to the field. Specifically, the use of artificial food substrates limited our evaluation of how larval predator avoidance (e.g., escape mechanisms) may alter competition outcomes in parasitized treatments. Other abiotic environmental conditions are also known to influence *L. heterotoma\* host selection (Roitberg et al. 1992). Unfavorable conditions such as changes in barometric pressure and humidity may diminish foraging strategies, leading to increased parasitization of suboptimal hosts in the field. Overall, complex and variable field conditions could potentially produce different ecological outcomes than those observed in this study.

## Conclusion

The enemy release hypothesis has raised controversy due to its mixed support within and across taxa (Colautti et al. 2004, Liu & Stiling 2006, Brian & Catford 2023). Our findings further emphasizes how varying contexts (e.g., density or competitor species) may alter species interactions and consequently affect the interpretation of enemy release. Here, we demonstrate that enemy release may contribute to the success of *Z. indianus*, and larval density influences the relative strength of enemy release. We confirm that *L. heterotoma* effectively decreases the emergence rates of *Z. indianus*, which may have major implications for management of *Z. indianus* in the field. However, these benefits may be diminished when more suitable hosts are present, especially at high larval densities. Due to the increasing threat of *Z. indianus* as an agricultural pest, productive biological control may control the growth and spread of *Z. indianus* populations. Verification of these results in a natural environment would be highly beneficial for understanding how this invasive species interacts with native ecosystems, and for determining if biological control could be an effective strategy for managing *Z. indianus* populations.

## Data Accessibility statement

All raw data used for these analyses and code to replicate the figures and statistics are available at https://doi.org/10.5061/dryad.6hdr7sr8n. Data and code are also available at https://github.com/camille-wa/Zaprionus-parasitoid-experiments.

## Author contributions

CRW and PAE conceptualized the study and acquired funding. CRW developed methodology, conducted all experiments, collected and analyzed all data, created figures, and wrote the manuscript with supervision from PAE. CRW and PAE revised the manuscript.

## Acknowledgements

We are grateful to Dr. Todd Schlenke for providing parasitoid wasp lines and rearing advice. We thank Adam Lenhart and Dr. Alan Bergland for *D. simulans* lines. We thank Logan Rakes and Weston Gray for assistance rearing *Z. indianus* lines. Dr. John Orrock and members of the Erickson lab provided helpful feedback on this project and manuscript. This work was funded by award # 1R15GM146208-01 to PAE from the National Institutes of Health, startup funds to PAE from the University of Richmond, a University of Richmond Summer Fellowship to CRW, and a Robert F. Smart Student Research Award to CRW.

## Appendix

**Appendix Table 1:** Generalized linear model results for each experiment. Z-values are reported for all experiments except for the interspecific competition assays (Fig. 1B-C), where T-values are given in italics due to use of quasibinomial errors. Significant terms were determined by P < 0.05 and are indicated in bold.

| Experiment | Term | Estimate | SE | Z/T | P-value |
| --- | --- | --- | --- | --- | --- |
| *Z. indianus* intraspecific<br>(Fig. 1A) | Intercept | 1.325 | 0.174 | 7.632 | < **0.001** |
|  | Parasitoid | -1.966 | 0.229 | -8.600 | < **0.001** |
|  | Density | -1.600 | 0.192 | -8.326 | < **0.001** |
|  | Parasitoid:Density | 1.230 | 0.260 | 4.729 | < **0.001** |
| *Z. indianus* vs *D. hydei* low density<br>(Fig. 1B-top) | Intercept | 0.717 | 0.364 | 1.971 | 0.066 |
|  | Parasitoid | -5.538 | 1.952 | -2.837 | **0.012** |
|  | Species | 0.350 | 0.535 | 0.654 | 0.522 |
|  | Parasitoid:Species | 3.084 | 2.036 | 1.515 | 0.149 |
| *Z. indianus* vs *D. hydei* high density<br>(Fig. 1B-bottom) | Intercept | -0.005 | 0.153 | -0.035 | 0.973 |
|  | Parasitoid | -3.404 | 0.460 | -7.402 | < **0.001** |
|  | Species | -1.316 | 0.242 | -5.445 | < **0.001** |
|  | Parasitoid:Species | 3.388 | 0.531 | 6.380 | < **0.001** |
| *Z. indianus* vs *D. simulans* low density<br>(Fig. 1C-top) | Intercept | 0.209 | 0.180 | 1.159 | 0.263 |
|  | Parasitoid | -5.029 | 1.021 | -4.924 | < **0.001** |
|  | Species | 0.736 | 0.269 | 2.737 | **0.015** |
|  | Parasitoid:Species | 3.256 | 1.059 | 3.076 | **0.007** |
| *Z. indianus* vs *D. simulans* high density<br>(Fig. 1C-bottom) | Intercept | -0.317 | 0.156 | -2.030 | 0.059 |
|  | Parasitoid | -4.911 | 1.071 | -4.586 | **0.003** |
|  | Species | 0.173 | 0.220 | 0.787 | 0.443 |
|  | Parasitoid:Species | 3.184 | 1.106 | 2.879 | **0.011** |
| *Z. indianus* Florida vs Connecticut<br>(Fig. 2) | Intercept | -2.753 | 0.400 | -6.890 | < **0.001** |
|  | State | -0.307 | 0.528 | -0.581 | 0.561 |
| *Z. indianus* vs. *D. simulans* exposure time<br>(Fig. 3) | Intercept | 0.547 | 0.219 | 2.499 | **0.013** |
|  | Species | 1.063 | 0.358 | 2.973 | **0.003** |
|  | 1 hour | -1.933 | 0.506 | -3.819 | < **0.001** |
|  | 8 hours | -3.914 | 1.040 | -3.762 | < **0.001** |
|  | 24 hours | -3.914 | 1.040 | -3.762 | < **0.001** |
|  | Species:1 hour | 1.171 | 0.704 | 1.664 | 0.096 |
|  | Species:8 hours | 1.899 | 1.141 | 1.665 | 0.096 |
|  | Species:24 hours | 2.573 | 1.139 | 2.258 | **0.024** |

